# An image-based CRISPR screen reveals splicing-mediated control of HP1α condensates

**DOI:** 10.1101/2025.09.21.676939

**Authors:** Matthew Man-Kin Wong, Shaopu Zhou, Chris Carpenter, Raeline Valbuena, Monika Priyadarshini, Aishwarya Arya, Aoon Rizvi, Catherine Carswell-Crumpton, Amelie Wileveau, Gabriel Lopez-Lopez, Josh Tycko, David Yao, Kaitlyn Spees, Jason C Maynard, Michael C Bassik, Hani Goodarzi, Serena Sanulli

## Abstract

Heterochromatin Protein 1α (HP1α) is a fundamental component of constitutive heterochromatin, forming subnuclear condensates whose regulation and function remain poorly understood. Here, we present an image-based CRISPR screen targeting nuclear factors that identifies splicing as a pivotal pathway regulating HP1α condensates. We discovered that unspliced intronic RNA modulates HP1α condensates by interacting co-transcriptionally with HP1α. By modulating the intron content, RNA processing restricts HP1α-RNA interactions at chromatin, thus enabling heterochromatin organization. Disruption of HP1α condensates due to enhanced interactions with unspliced RNA leads to loss of heterochromatin and the activation of stress response protective genes. We propose that RNA is a central component of heterochromatin that modulates HP1α condensates, and that RNA processing enzymes act as a surveillance mechanism for condensates by dynamically regulating the network of multi-valent interactions between RNA and chromatin factors. This model underscores the crosstalk between chromatin organization, transcription, and RNA processing, potentially governing broader nuclear functions.

---

Constitutive heterochromatin is a highly conserved and structurally distinct portion of the eukaryotic genome, essential for genome stability and transcriptional silencing^1^. This complex platform consists of multiple core components, with Heterochromatin Protein 1α (HP1α) at its center. HP1α acts as a structural adaptor that recognizes the methylated lysine 9 of histone H3 (H3K9me), compacts chromatin, and recruits heterochromatic-associated factors^2^. In cells, HP1α forms discrete subcellular structures, appearing as dynamic spherical foci of about 1 μm in diameter, that can fuse into larger bodies and dissolve during mitosis^3,4^. Within the foci, HP1α molecules are highly concentrated yet diffuse rapidly, exchanging within seconds^4–6^. Due to the dynamic nature and biophysical properties of HP1α, phase separation has been implicated in the formation of HP1α foci by creating membraneless compartments in which chromatin is compacted, and heterochromatin factors are enriched^4,7,8^.

RNA is also a crucial but less understood component of mammalian heterochromatin^9^. Although the presence of RNA within silenced genomic regions might seem counterintuitive, several lines of evidence indicate its important role. In plants, worms, and yeast, it is well established that RNA interference (RNAi) pathways are involved in the formation and maintenance of pericentric heterochromatin^1,9^. In mammals, an RNAi-dependent mechanism has not been described, but RNA appears to serve a structural function. Depleting RNA in cells with RNAse treatment results in the loss of HP1α foci, and defects in the compaction and structural integrity of heterochromatin^3,10^. Furthermore, in mouse embryonic stem cells, RNA from satellite repeats is crucial for recruiting HP1α at pericentromeric regions^11^. How RNA regulates heterochromatin in mammals and whether it directly interacts with HP1α in cells is unclear.

The molecular mechanisms underlying heterochromatin organization and the clustering of HP1α molecules in condensates also remain poorly understood. Efforts to elucidate these mechanisms have been hindered by the lack of high throughput image-based approaches to visualize such structures. Here, we aimed to investigate the cellular pathways that regulate the localization of HP1α in nuclear foci, with the goal of uncovering the principles that govern the spatial and functional organization of heterochromatin. To overcome current technical limitation, we developed an image-based CRISPR screen for regulators of HP1α condensates. Through this approach, we discovered that HP1α condensates are modulated by direct interaction with intronic RNA. Our findings provide an opportunity to revise current models for heterochromatin by describing a regulatory crosstalk between RNA processing and heterochromatin.

Here, we use the term “condensate” to denote subcellular structures that appear as microscopic foci with high local concentrations of specific molecules, without making any assumptions about their physical properties or material states.

## Development of a CRISPR KO screen for HP1α condensates

To dissect the mechanisms driving HP1α organization in condensates, we developed a pooled image-based CRISPR knockout (KO) screen for regulators of HP1α nuclear distribution. We reasoned that any perturbation altering the interactions underlying HP1α condensates would affect the nuclear distribution of HP1α, leading to the loss of foci or changes in their intensity and size.

We infected K562 human cells expressing Cas9 with a custom lentiviral CRISPR KO library of single-guide RNAs (sgRNAs) targeting 5,373 nuclear factors (Fig. 1a), then fixed and stained cells with an HP1α primary antibody and a fluorescent secondary antibody to detect endogenous protein (Fig. 1b). We separated cells based on HP1α signals using high-speed image-enabled cell sorting (ICS) (Fig. 1a, Extended Data Fig. 1a). Unlike conventional cell sorting, ICS can quantify the subcellular localization of labeled proteins and perform image-derived sorting decisions following real-time image analysis^12^. To identify the parameters that best differentiate changes in foci, we compared K562 wild-type (WT) cells, *HP1α* KO cells (Fig. 1b), and cells with the sgRNA library. The combination of two parameters best captured the features of HP1α subcellular localization: maximum intensity, which reflects the brightest pixel value within each cell, and signal size, defined by the number of pixels above a user-defined signal threshold (Extended Data Fig. 1b). Importantly, these two features describe the overall distribution of HP1α signal in cells and cannot be simply interpreted as an increase in HP1α foci size and intensity (Fig. 1c). Leveraging these two features, we established a hierarchical gating strategy for cell sorting and defined two cell populations, which we called “high” and “low”, that significantly deviated from WT cells (Fig. 1c). We subsequently isolated the genomic DNA, measured the sgRNA frequencies in the two populations via deep sequencing, and analyzed our results using the Cas9 high-throughput maximum likelihood estimator (CasTLE) algorithm, which provides a confidence score for the effect of each gene knockout^13^.

**Figure 1:**
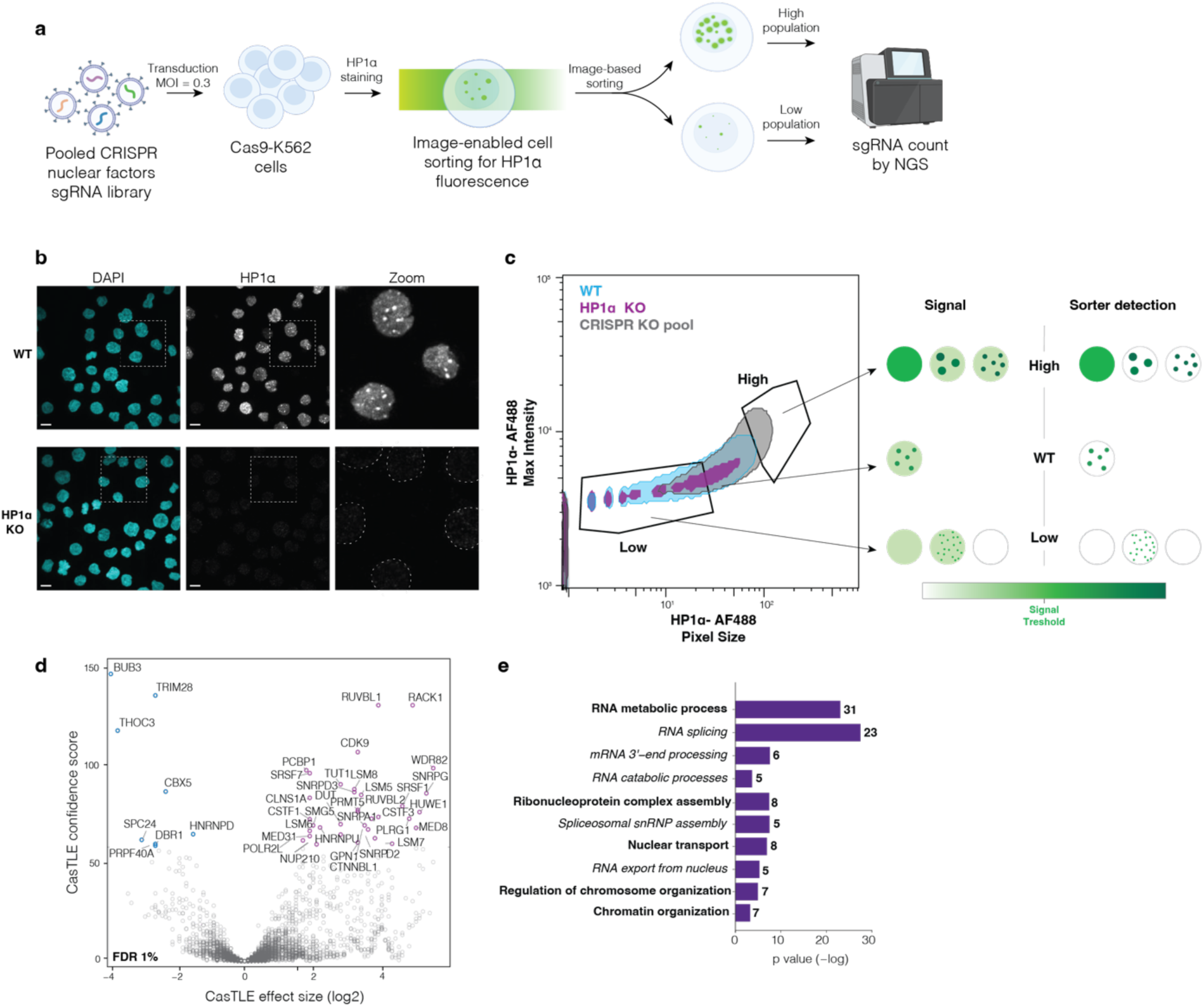
Image-based screen for HP1α-condensate regulators. **a,** Image-based pooled KO CRISPR screen targeting nuclear factors in K562 cells. Following library installation, cells are immunostained for endogenous HP1α and sorted into “high” and “low” populations based on the Alexa Fluor 488 dye (AF-488)-HP1α signal. Genomic extraction and analysis are performed to identify sgRNA enriched in the two populations. **b,** K562 WT and *HP1α* KO cells stained for HP1α (gray). Nuclei are stained with DAPI (cyan). Scale bar: 20 μm. **c,** Contour plot of WT (blue), HP1α KO (purple), and CRISPR KO (gray) cells based on maximum intensity (y-axis) and pixel size (x-axis) of the AF-488-HP1α signal. A schematic of potential HP1α signals in the different populations is shown on the right. **d,** Volcano plot of the screen described in a. Dotted line indicates 1% FDR. All 42 hits are shown. Blue and purple circles represent hits with negative and positive CasTLE scores, respectively. **e,** GO enrichment analysis based on biological functions for hits with a significant effect size shown in d. Gene classes are in bold and subclasses in italics. Bold numbers indicate the number of genes per class. MOI: multiplicity of infection; NGS: next-generation sequencing.

Using a false discovery rate (FDR) of 1%, we identified 42 genes predicted to regulate HP1α localization in foci (Fig. 1d, Extended Data Fig. 1c-e). Eight of these genes are enriched in the low population (negative CasTLE effect), while the remaining 34 genes are enriched in the high population (positive CasTLE effect). *CBX5*, which encodes HP1α, is among the top screen hits with a negative CasTLE effect, validating the image-based screening platform. Remarkably, except for Tripartite motif-containing 28 (*TRIM28*), which codes for an HP1α-interacting factor^14^, none of the remaining hits are known components of the heterochromatin pathways. Instead, gene ontology (GO) analysis indicates that over 70% of the genes (31/42 genes) are involved in RNA metabolic processes, specifically RNA splicing, RNA catabolic processes, and messenger RNA (mRNA) 3’ processing (Fig. 1e, Extended Data Fig. 1e). The remaining genes are involved in (i) ribonucleoprotein complex assembly, (ii) nuclear transport and RNA export from the nucleus, (iii) regulation of chromosome segregation, and (iv) chromatin organization.

Overall, this new image-based screen for regulators of HP1α condensates uncovered an unexpected link between HP1α localization and RNA processing.

## Correlation between increased intron retention and HP1α mislocalization

To validate the screen hits and further investigate the link between HP1α and RNA processing, we generated single-KO cell lines of the top hits and determined the localization of endogenous HP1α protein by confocal imaging (Fig. 2a). We selected genes that (i) presented high-confidence CasTLE scores, (ii) were anticipated to be non-essential for cell viability, and (iii) are involved in different steps of RNA processing. To control for compensatory and indirect effects, KO cell lines were generated as a pool via lentivirus infection and used at low passage (<5) and KO efficiency confirmed by sequencing based on inference of CRISPR edits (ICE)^15^ (Extended Data Fig. 2a). Based on these criteria, we focused on the following genes: *HP1*α, *TRIM28*, THO complex subunit 3 (*THOC3*), cleavage stimulation factor subunit 3 (*CSTF3*), GPN-loop GTPase 1 (*GPN1*), and WD repeat domain 82 (*WDR82*).

**Figure 2:**
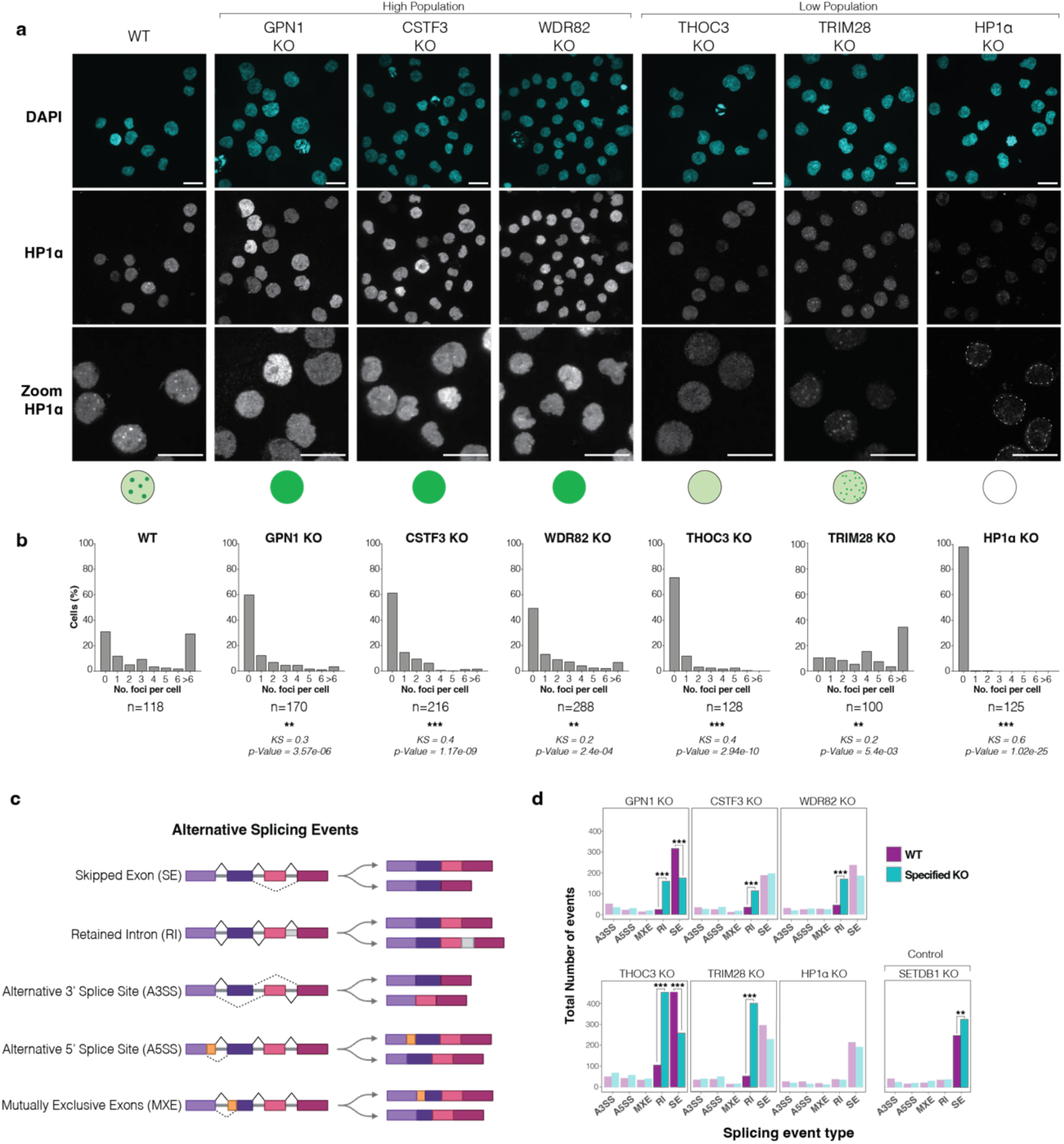
Mutants with altered HP1α foci present increased IR. **a,** Maximum intensity projection confocal images of K562 WT and KO cells immunostained for HP1α (gray). Nuclei are stained with DAPI (cyan). Magnified regions are shown in the lower panels, with schematic illustrations of the observed phenotypes. Scale bar: 20 μm. **b,** Quantification of HP1α foci per cell. *n*: number of cells analyzed. Kolmogorov-Smirnov (KS) statistics and corresponding *p*-values compare the distribution of each KO to WT cells. **c,** Schematic illustrating the types of AS events analyzed in panel d. **d,** Total number of AS events in KO vs. WT cells. Bars representing non-significant changes are shown in transparent colors for clarity (*p* ≤ 0.01). Black asterisks indicate events significantly different in the KO.

TRIM28, aside from being a component of heterochromatin, also regulates RNA polymerase II (Pol II) pausing and transcriptional elongation^14^. *THOC3* is important for efficient export of polyadenylated and spliced RNA^16^. It encodes a component of the nuclear THO transcription elongation complex, which binds spliced mRNA and facilitates its export to the cytoplasm^17^. *CSTF3* encodes a subunit of the CSTF complex, which is involved in the polyadenylation and 3’-end cleavage of pre-mRNAs^18^. *GPN1* encodes a guanosine triphosphatase enzyme that is required for the proper assembly and nuclear import of RNA Pol II, acting as a bridge between RNA Pol II and scaffolding proteins^19^. WDR82 plays a key role in tethering the histone H3-Lys4 methyltransferase complex SET1 to transcriptional start sites and facilitating its interaction with the Ser-5’ phosphorylated C-terminal domain of RNA Pol II^20^.

We performed quantitative and statistical analysis of endogenous HP1α localization using the image analysis pipeline Punctatools^21^ on three-dimensional z-stack confocal images (Fig. 2a, b, Extended Data Fig. 2b). We analyzed a minimum of 100 cells per condition to ensure robust representation of the phenotype.

While all KO cells maintain nuclear HP1α localization, they display a range of phenotypes, including changes in the number of puncta per cell, fluorescence intensity, and puncta volume. Importantly, we observe that (i) different HP1α nuclear distributions can be represented by different combinations of signal intensity and pixel size (Fig. 2a, Extended Data Fig. 2c, d), and (ii) due to the complexity of HP1α signal changes, CasTLE scores reflect a range of complex phenotypes that cannot be read as simply positive or negative regulators of HP1α foci, as described below.

In WT cells infected with a control safe gRNA that targets a non-essential genomic region, HP1α exhibits a heterogeneous distribution of foci count per cell: approximately 30% of cells show no foci, 70% present between 1-5 foci per cell, and 30% display more than 6 foci (Fig. 2a, b). This finding is consistent with previous observations of chromocenter organization in mammalian cells^22^. As expected, *HP1α* KO cells present no foci (Fig. 2a, b). *GPN1*, *CSTF3*, *WDR82*, and *THOC3* KO cells show a reduced number of HP1α foci per cell, with the majority of nuclei (>60%) presenting none (Fig. 2a, b). *TRIM28* KO cells present an overall increase in the number of foci per cell: only 10% of cells present no foci, with the majority (63%) presenting 4 foci or more (Fig. 2a, b). In *TRIM28* KO the foci present a slight decrease in volume and overall lower intensity compared with WT cells (Extended Data Fig. 2c, d), which may be linked to changes in chromocenter organization.

Overall, all mutants from both the high and low populations show changes in the HP1α subcellular distribution, indicating that the screen was effective in discovering regulators of HP1α condensates. KO mutants enriched in the high population (*GPN1*, *CSTF3*, and *WDR82* KO) present a general loss of foci and increased HP1α nucleoplasm staining. This phenotype was decoded by the sorter as an increase in the positive pixels size and intensity per cell above a set threshold intensity (Fig. 1c). KO mutants enriched in the low population (*THOC3*, *TRIM28* KOs) present different phenotypes: *TRIM28* KO cells have more puncta at lower intensity and volume, whereas *THOC3* KO cells present a general loss of foci. In both cases, the sorter interpreted the phenotype as a decrease in the positive pixels size and intensity per cell relative to the set threshold intensity. Hereafter, we refer to these KO cells as “HP1α localization mutants”.

To ensure that HP1α mislocalization is not merely a reflection of altered HP1α expression levels, we compared both transcript and protein abundances across the KO cells. No significant changes between mutants and WT cells are observed at the RNA level (Extended Data Fig. 2e). Although *CSTF3* and *WDR82* KO cells show a slight increase in HP1α protein level, the remaining mutants maintain HP1α levels comparable to those in WT cells (Extended Data Fig. 2f). Thus, we reason that the increase in the HP1α nucleoplasmic signal observed by immunostaining is likely due to the dispersion of HP1α from highly concentrated foci into the whole nucleoplasm and/or an alteration in epitope accessibility^23^. The localization of HP1β, a paralog of HP1α, appears unchanged across these mutants (Extended Data Fig. 2g).

Given that RNA splicing is the most enriched GO class among our screening hits, we hypothesized that alterations in RNA splicing might be occurring in the HP1α localization mutants. Thus, we performed RNA sequencing (RNA-seq) and analyzed alternative splicing events across all the HP1α localization mutants (Fig. 2c). KO cells for the H3K9 methyltransferase *SETDB1* were included as a negative control, as they present no changes in HP1α foci (Extended Data Fig. 2h). All five HP1α localization mutants display a significant increase in intron retention (IR) events (Fig. 2d, Extended Data Fig.2i, j). IR was further validated using IRFinder (Extended Data Fig. 2k), a tool tailored to IR analysis^24^, and long-read RNA sequencing (Extended Data Fig. 2l, m). No significant IR changes are observed in *HP1α* or *SETDB1* KO cells (Fig. 2d, Extended Data Fig. 2k), indicating that the loss of HP1α or heterochromatin defects do not directly regulate IR.

Alternative splicing is a co-transcriptional regulatory mechanism that can affect cellular functions by modulating transcriptome and proteome diversity^25,26^. To elucidate the relationship between IR and HP1α condensates, we investigated whether IR events occur within genes that are either differentially expressed or enriched for specific pathways. All KO mutants, with the exception of *TRIM28*, exhibit minimal transcriptome-wide changes when compared to WT (Extended Data Fig. 2n, o). mRNA transcripts with IR are neither differentially expressed nor associated with any common gene categories (Extended Data Fig. 2n-p), suggesting that the changes in HP1α condensates are unlikely to be driven by alterations in gene expression. In contrast, the widespread upregulation in gene expression observed in *TRIM28* KO cells (Extended Data Fig. 2n, o) aligns with TRIM28’s known roles in both RNA processing and gene silencing^14^. To assess the role of AS in regulating HP1α condensates independently of transcriptional changes, we focused on characterizing the localization mutants that do not display global gene expression alterations, namely *THOC3, CSTF3, GPN1,* and *WDR82* KOs.

Next, to investigate whether AS regulates HP1α interactors required for condensate formation, we performed HP1α Immunoprecipitation followed by mass spectrometry (IP-MS) in WT, *CSTF3* KO, and *THOC3* KO cells. No common HP1α interactor undergoes AS across all localization mutants (Extended Data Fig. 2p), and only a small subset of distinct factors (none in *CSTF3* KO and 10/314 for *THOC3* KO) show both differential AS and reduced interaction with HP1α (Extended Data Fig. 2p, r). Importantly, no changes in the splicing isoforms of HP1α itself are detected in the localization mutants, indicating that the mislocalization is not linked to altered splicing of HP1α (Extended Data Fig. 2s).

In summary, *TRIM28, THOC3, CSTF3, GPN1,* and *WDR82* are *bona fide* regulators of HP1α condensates in cells. The increase in IR across all localization mutants suggests a link between intronic RNA and condensate formation. Genes exhibiting IR are (i) not differentially expressed, (ii) not common across the different KOs, (iii) not linked to a common biological pathway, and (iv) not affecting a common HP1α interactor. We conclude that a post-transcriptional regulatory mechanism underlies the changes in HP1α foci in the localization mutants.

## HP1α binds introns within heterochromatin and shows phase re-entry behavior

RNA is important for heterochromatin organization and for HP1α condensate formation^3,10,11^. HP1α contains a nucleic-acid-binding domain, called the hinge, which is unstructured, positively charged, and capable of binding both RNA and DNA *in vitro* (Fig. 3a)^27–29^. We hypothesized that the correlation between IR and changes in HP1α localization might be due to altered direct interaction between RNA and HP1α.

**Figure 3:**
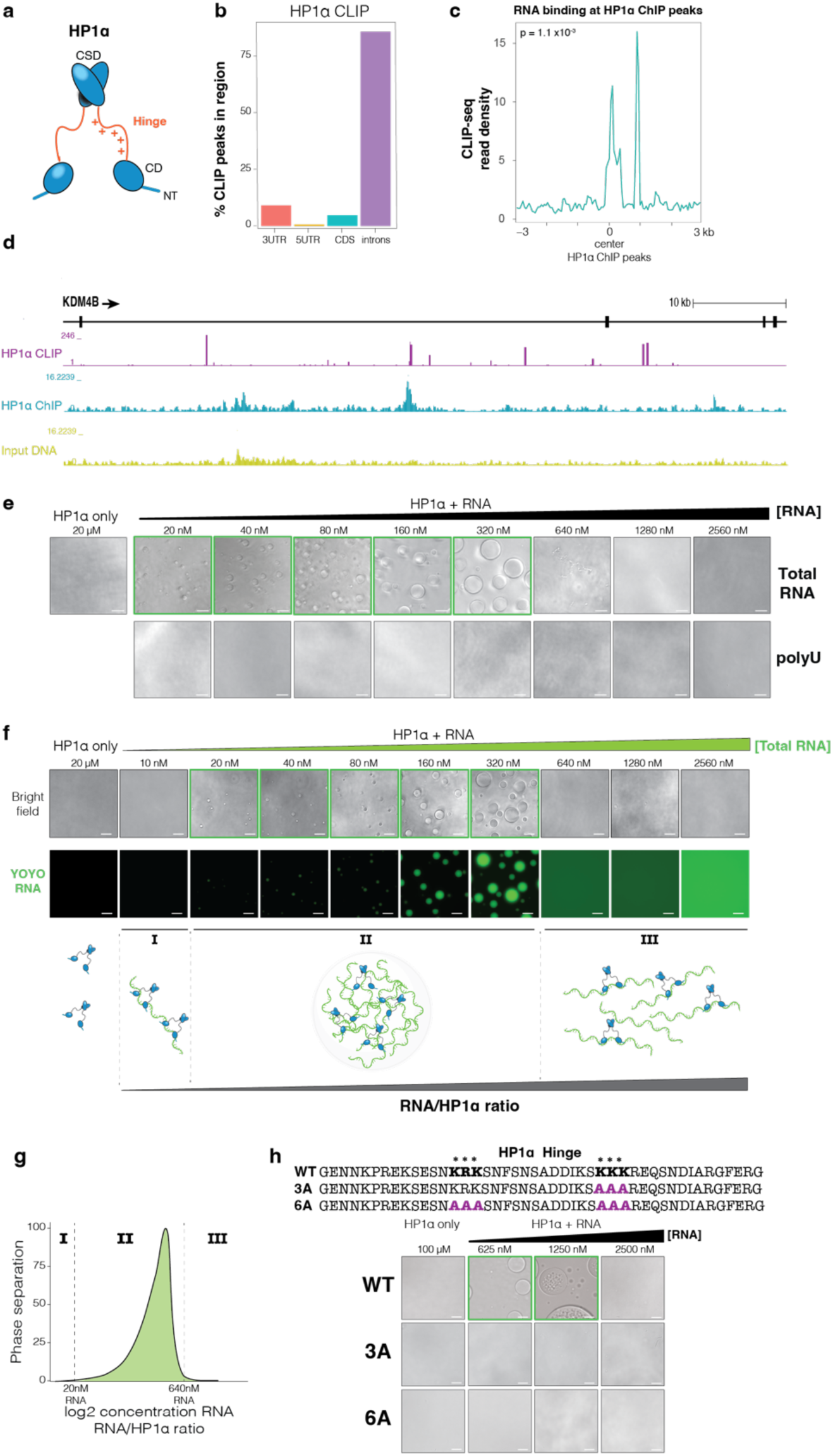
HP1α is an RNA-binding protein that interacts with introns. **a,** Schematic of HP1α protein domains. The Hinge domain is disordered and positively charged. CSD: chromoshadow domain; CD: chromodomain; NT: N-terminal domain. **b,** Relative distribution of HP1α CLIP-seq peaks across transcript features in K562 WT cells. CDS: coding sequence. **c,** Metagene plot showing enrichment of HP1α–RNA binding at genomic intronic regions associated with HP1α. *p*-value from Pearson’s Chi-Square test of independence is indicated. **d,** Genome browser view showing both HP1α CLIP- and ChIP-seq peaks at the *KDM4B* locus. **e,** Bright-field images of hHP1α (20 μM) phase separation assays with increasing concentration of total or polyU RNA (20-2560nM). Scale bar: 4 μm. **f,** Bright-field and fluorescence microscopy images of phase separation assays as in panel e, using YOYO-labeled RNA. RNA concentrations range 10-2560nM, HP1α is 20μM. Scale bar: 4 μm. A cartoon below illustrates molecular interactions across three regimes (I-III), showing homogenous solution in I and III, and phase separation in II. **g,** Phase diagram quantifying RNA-induced phase separation from panel f. **h**, Top: schematic of the 3A and 6A alanine substitutions within the Hinge domain. Bottom: bright-field images of phase separation assays using WT, 3A or 6A HP1α (100μM) with increasing RNA concentrations (0.625-2.5mM). Scale bar: 4 μm.

Before exploring this possibility, we first evaluated HP1α-RNA interactions in WT cells under normal steady-state conditions by performing HP1α cross-linking and immunoprecipitation sequencing (CLIP-seq). The majority (82.1%) of HP1α CLIP signal is within protein coding RNAs (Extended Data Fig. 3a). Remarkably, over 60% of peaks localize to introns (Fig. 3b) of transcripts with low GC content (40%) and involved in a wide range of biological processes (Extended Data Fig. 3b-d). HP1α exhibits widespread, low-specificity binding to low-complexity RNA motifs, suggesting context-dependent interactions or biological heterogeneity in its binding sites (Extended Data Fig. 3e).

It is well established that introns are co-transcriptionally spliced^30–33^. Therefore, we reasoned that HP1α might interact with unspliced RNA in proximity of its chromatin-binding regions before intron removal by splicing. Re-examining previously published chromatin immunoprecipitation (ChIP) data, we observed that HP1α peaks (36.5%) are significantly enriched in intronic genomic regions (*p* = 0.017, Z-score = 2.118) (Extended Data Fig. 3f). Metaplot analysis shows that the HP1α CLIP signal is statistically enriched at HP1α ChIP sequencing (ChIP-seq) peaks corresponding to introns (*p* = 0.002, Z-score = 3.53), indicating that HP1α binds introns in both genomic DNA and RNA (Fig. 3c). Overall, CLIP peaks are found both in correspondence of ChIP peaks and of other nearby intronic sites, as shown for the *KDM4B* locus (Fig. 3d). This spatial coupling between RNA binding and chromatin association suggests that at least a subset of RNA-HP1α interactions may occur co-transcriptionally at or near its chromatin-associated sites; however, distinguishing *cis* from *trans* contributions will require further investigation.

Given that HP1α is a marker for heterochromatic regions typically considered silenced, we sought to further assess the chromatin and transcriptional landscape at HP1α genomic locations. As expected, H3K9me3 is enriched at HP1α ChIP peaks (Extended Data Fig. 3g). We then examined the expression levels of the genes and transcripts that are bound by HP1α. Consistent with previous studies, the distribution of the whole transcriptome shows a primary peak of highly expressed genes, with a left shoulder of lowly expressed transcripts^34^ (Extended Data Fig. 3h). Surprisingly, a subset of transcripts corresponding to all HP1α-bound genes, and genes that are both HP1α-bound and H3K9me-enriched display a similar distribution, indicating that transcription occurs within regions commonly defined as heterochromatic (Extended Data Fig. 3h). The subset of transcripts with HP1α CLIP peaks is more enriched for highly expressed RNAs (Extended Data Fig. 3h), which might indicate that HP1α preferentially binds RNA from highly expressed gene or could reflect a technical limitation in detecting low levels of intron-containing transcripts. Altogether, these results indicate that transcription can occur within heterochromatic regions marked by HP1α and H3K9me3, and that HP1α binds to RNA produced near its genomic binding sites.

To further dissect the HP1α–RNA interaction, we performed *in vitro* phase separation assays with recombinant human HP1α (hHP1α) and different types of RNAs. We employed either polyuridylic acid (polyU) RNA, consisting of molecules with a length range of 600–3000 nucleotides (nt), or total cellular RNA containing molecules of various sizes. In this assay, we varied the RNA concentration monotonically against a constant hHP1α concentration and visualized droplets via phase contrast microscopy. PolyU RNA induces phase separation at high micromolar concentrations of HP1α and RNA^11^(Extended Data Fig. 3i), but no phase separation is observed at HP1α concentrations that mimic endogenous cellular levels^35,36^ (Fig. 3e); on the contrary, total cell RNA promotes phase separation at physiological concentrations of hHP1α (Fig. 3e). Using 3% YOYO-labeled RNA, we observed that HP1α undergoes sequential demixing and mixing phase transitions in response to increasing RNA concentration, with RNA becoming enriched inside the condensates (Fig. 3f). We constructed a phase diagram based on fluorescent microscopy and identified three distinct regimes defined by upper and lower critical RNA concentrations, C_1_ and C_2_. At concentrations below C_1_ (20nM) or above C_2_ (350 nM), one homogeneous phase prevails. At RNA concentrations between C_1_ and C_2_, two liquid phases coexist, with droplets size increasing as RNA concentration rises (Fig. 3f, g). This phenomenon is consistent with previously described phase re-entry behavior in the context of RNA and RNA-binding proteins^37^, where the RNA promotes both droplet assembly and dissolution depending on the RNA–protein ratio. At lower RNA–HP1α ratios, RNA nucleates condensate formation, whereas at higher ratios, RNA leads to dissolution.

The HP1α Hinge domain is known to mediate both RNA and DNA binding *in vitro*^29,38,39^. Mutants in which three (3A) or six (6A) positively charged residues within the Hinge are replaced with alanine fail to undergo RNA-mediated phase separation *in vitro*, confirming that this domain is essential for RNA-driven condensate formation (Fig. 3h, Extended Data Fig. 3j).

Together, these results demonstrate that HP1α is an RNA-binding protein that binds GC-poor sequences within introns of transcripts implicated in a wide range of cellular functions. Since RNA splicing occurs co-transcriptionally and HP1α interacts with RNAs transcribed near its chromatin-binding site, we conclude that HP1α–RNA interactions preferentially occur at chromatin. This observation, consistent with recent studies reporting transcription within regions traditionally considered silent^40^, suggests that transcription can occur within heterochromatic regions.

## RNA both promotes and inhibits HP1α-chromatin condensates

HP1α can bind to and phase separate with both RNA and H3K9me3 chromatin individually (Fig. 3e,f and Extended Data Fig. 4a-c)^7,8,11^, indicating that HP1α can establish heterotypic interactions with both substrates. Given that the Hinge domain participates in both interactions, and that RNA and chromatin coexist in the nucleus, we explored whether RNA and chromatin act synergistically or compete for HP1α binding.

We characterized the phase behavior of HP1α with varying concentrations of RNA and H3K9me3 chromatin. At low concentrations, RNA promotes phase separation by reducing the chromatin critical concentration (C_c_) required for condensate formation (Fig. 4a and Extended Data Fig. 4d), indicating cooperative binding between RNA and chromatin when HP1α concentration is not limiting. The finding that RNA promotes HP1α condensation with H3K9me chromatin is consistent with the requirement of RNA for HP1α foci formation in cells^3,10,11^. In contrast, at high concentrations, RNA inhibits phase separation (Fig. 4a, b and Extended Data Fig. 4d), suggesting competition between RNA and chromatin when HP1α amount becomes limiting.

**Figure 4:**
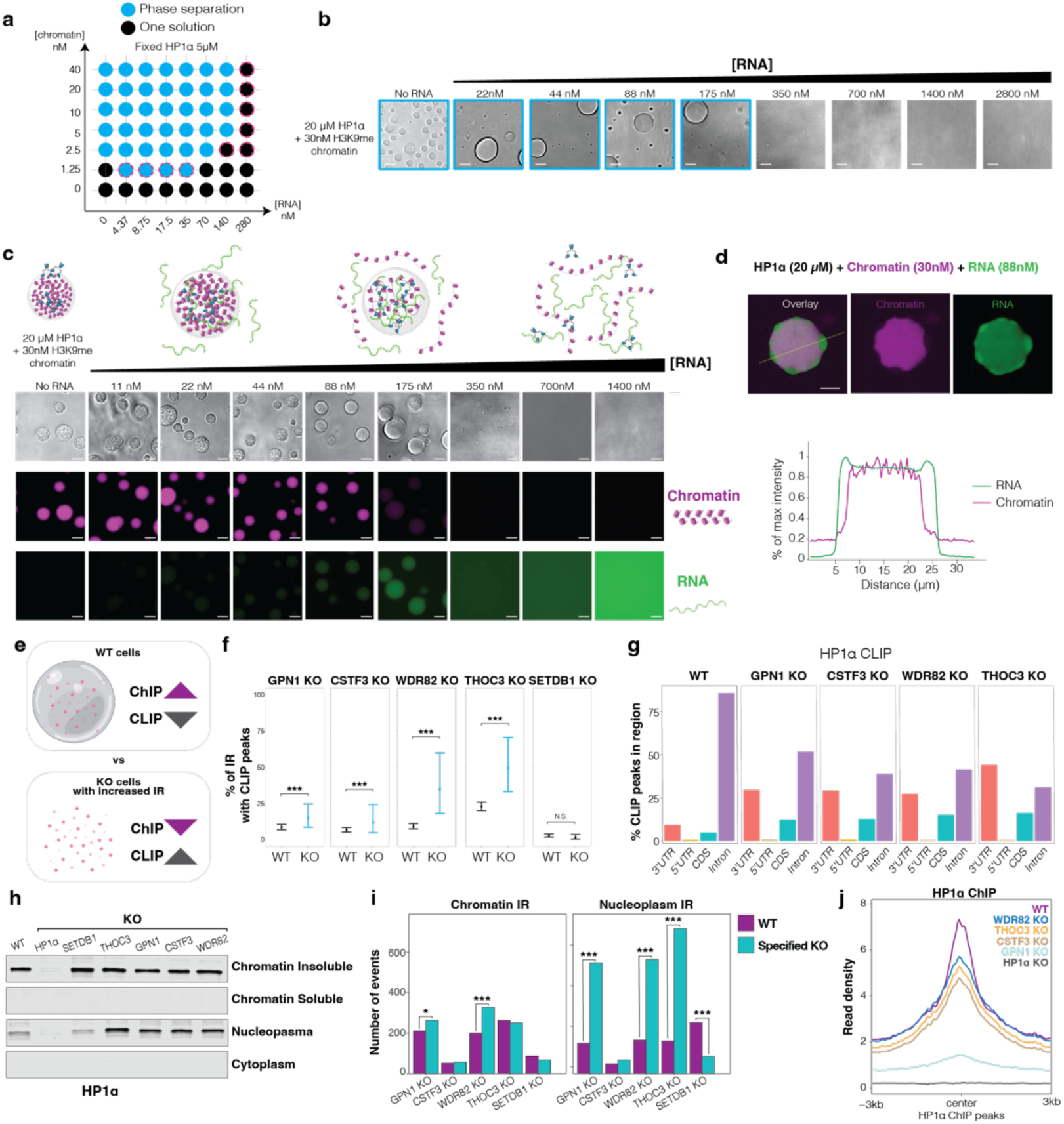
RNA modulates HP1α–chromatin condensates. **a,** Phase diagram showing phase separation as a function of RNA and chromatin concentration at a fixed hHP1α concentration (5μM). Blue circles indicate conditions where phase separation occurs; black circles indicate conditions without phase separation. Dashed magenta circles highlight conditions where RNA alters the C_c_. Corresponding microcopy images are shown in Extended Data Fig. 4d. **b,** Bright-field images of phase separation assays with increasing concentrations of total cellular RNA and fixed concentrations of hHP1α (20 μM) and H3K9me3 chromatin (30nM). RNA concentrations are serial 1:2 dilutions starting from 2.8 mM. Blue squares mark conditions where phase separation is observed. Scale bar: 4 μm. **c,** Bright-field and fluorescent microscopy images of phase separation assays as in panel b, using 3% Cy-3-labeled H3K9me3 chromatin and 3% YOYO-labeled total RNA. Scale bar: 4 μm. A schematic on top illustrates molecular interactions. **d,** Top: Confocal section of an *in vitro* reconstituted three-component condensate with hHP1α, YOYO-labeled RNA, and Cy-3-labeled H3K9me3 chromatin. Bottom: Line scan showing fluorescence intensity profiles of chromatin and RNA across the yellow line. Scale bar: 10 μm. **e,** Model for condensate regulation. **f,** Percentage of retained introns containing HP1α CLIP peaks in localization mutants and WT cells. *SETDB1* KO serves as a negative control. **g,** Distribution of HP1α CLIP peaks across transcript features in localization mutants. CDS: coding sequence. **h,** WB of HP1α in distinct cellular fractions from WT and KO cells. **i,** Quantification of IR in chromatin-associated and nucleoplasmic RNA fractions in WT and KO cells. **j,** Metagene plots of HP1α ChIP-seq signal in KO cells compared to WT. HP1α KO cells serve as negative control.

To further investigate the competition model, we assessed RNA and chromatin partitioning using YOYO-labeled-RNA and Cy5-labeled H3K9me3 chromatin. At the lowest RNA concentration, both RNA and chromatin co-partition in the droplets. As the RNA concentration increases, RNA partitioning in the droplets increases while chromatin enrichment decreases (Fig. 4c). This result indicates that RNA can enter the HP1α–chromatin condensates and displace chromatin in a concentration-dependent manner. The HP1α–chromatin droplets displayed visible internal heterogeneity (Fig. 4c) suggesting the presence of sub-structures with different molecular compositions. Therefore, to evaluate the spatial organization of RNA and chromatin within phase-separated droplets, we used confocal microscopy to image *in vitro* reconstituted three-component condensates. We observed that chromatin is enriched at the core of the droplet, while RNA accumulates in surface “vacuoles” or voids where chromatin is depleted (Fig. 4d). Similar multiphase structures have been observed in various biological contexts involving multi-component condensates and are thought to arise from competition for a shared binding partner^41,42^.

To further explore this competitive interaction, we incubated H3K9me3 chromatin with increasing concentrations of the HP1α Hinge mutants 3A and 6A. Both mutants show impaired phase separation with chromatin (Extended Data Fig. 4e), supporting a model in which chromatin and RNA compete for binding to the HP1α Hinge domain. This finding is consistent with previous studies indicating that the Hinge domain contributes to chromatin binding through interaction with nucleosomal DNA^29^.

Given HP1α’s interaction with intronic RNA and the observed competition between RNA and chromatin, we hypothesize that transcripts with retained introns may regulate HP1α condensates in the localization mutants. If intron-retaining transcripts relocalize HP1α through direct binding, we would expect to observe enhanced HP1α–RNA interactions and reduced HP1α–chromatin interactions in these mutants (Fig, 4e). CLIP-seq analysis in all localization mutants revealed increased HP1α binding to retained introns compared to WT cells (Fig. 4f, Extended Data Fig. 4f). Similar to WT, the HP1α-bound introns in the mutants tend to have lower GC content and average length of 500 nt (Extended Data Fig. 4g, h). Notably, analysis of CLIP peak distribution revealed that in the localization mutants, HP1α shows an increase in peaks at non-intronic regions, particularly within the 3’ untranslated regions (UTRs) and coding sequences (CDSs) (Fig. 4g). This suggests that HP1α not only enhances its interactions with introns but also gains binding to additional regions within the same transcripts (Extended Data Fig. 4i). This shift may reflect HP1α binding to transcripts that are no longer chromatin-associated and have undergone splicing, resulting in an overall reduction in intron content.

To assess HP1α chromatin association, we performed biochemical fractionation and observed increased HP1α in the nucleoplasmic fraction across all localization mutants (Fig. 4h, Extended Data Fig. 4j). To investigate where retained introns accumulate, we fractionated cellular RNA and quantified introns in both chromatin and nucleoplasmic fractions^43^ (Extended Data Fig. 4k). As expected, intron-containing RNAs are primarily chromatin-associated (Extended Data Fig. 4l). However, we found an accumulation of intronic RNAs in the nucleoplasm across all localization mutants (Fig. 4i, Extended Data Fig. 4m). Combined with the increase of HP1α in the nucleoplasm, this suggests weakened chromatin association of HP1α. To further confirm this, we performed spike-in ChIP-seq in the localization mutants (Fig. 4j) and observed reduced HP1α-chromatin association together with increased RNA CLIP signal at these sites (Extended Data Fig. 4n), indicating a shift from chromatin-bound to RNA-associated HP1α.

Finally, to test the functional role of the Hinge domain in HP1α localization, we performed ChIP-seq in *HP1α KO* cells rescued with 3A and 6A hinge mutants (Extended Data Fig. 4o). Both mutants showed reduction in chromatin association and foci (Extended Data Fig. 4p, q), supporting the importance of nucleic acid binding for HP1α localization.

Taken together, our findings demonstrate that RNA regulates HP1α condensates *in vitro* by directly interacting with HP1α, and that increased HP1α-RNA interactions relocalize HP1α in cells. We propose that chromatin-associated RNA within heterochromatin facilitates HP1α condensate formation with H3K9me by increasing HP1α local concentration on chromatin. When intron-containing transcripts accumulate in the nucleoplasm, they compete with chromatin for HP1α binding, leading to its relocalization and loss of condensates.

## IR upon stress disrupts HP1α condensates to activate protective genes

Intron-containing transcripts are typically chromatin-associated, lowly abundant, and transient due to co-transcriptional splicing^30–33^. Under certain physiological conditions, such as heat stress, IR increases, leading to the accumulation of intron-containing transcripts in the nucleoplasm^44–46^. This phenomenon has been proposed to act as a post-transcriptional regulatory mechanism that helps cells adapt to and recover from stress^44,45^. The precise molecular mechanisms by which IR regulates the stress response remain unclear.

To evaluate whether increased IR regulates HP1α condensates in physiological conditions, we induced heat stress and monitored HP1α localization. AS analysis over a time course reveals that IR is greatest at 2 hours of heat stress (Extended Data Fig. 5a). A general loss of HP1α condensates during stress is observed, indicating that the link between IR and HP1α localization is preserved in physiological responses (Fig. 5a, b, Extended Data Fig. 5b). Notably, HP1β localization remains unchanged under these conditions (Extended Data Fig. 5c). A similar loss of HP1α condensates is also observed following treatment with the splicing inhibitors Pladienolide B, further indicating that splicing perturbations – regardless of their origin – can alter HP1α condensates (Extended Data Fig. 5d).

**Figure 5:**
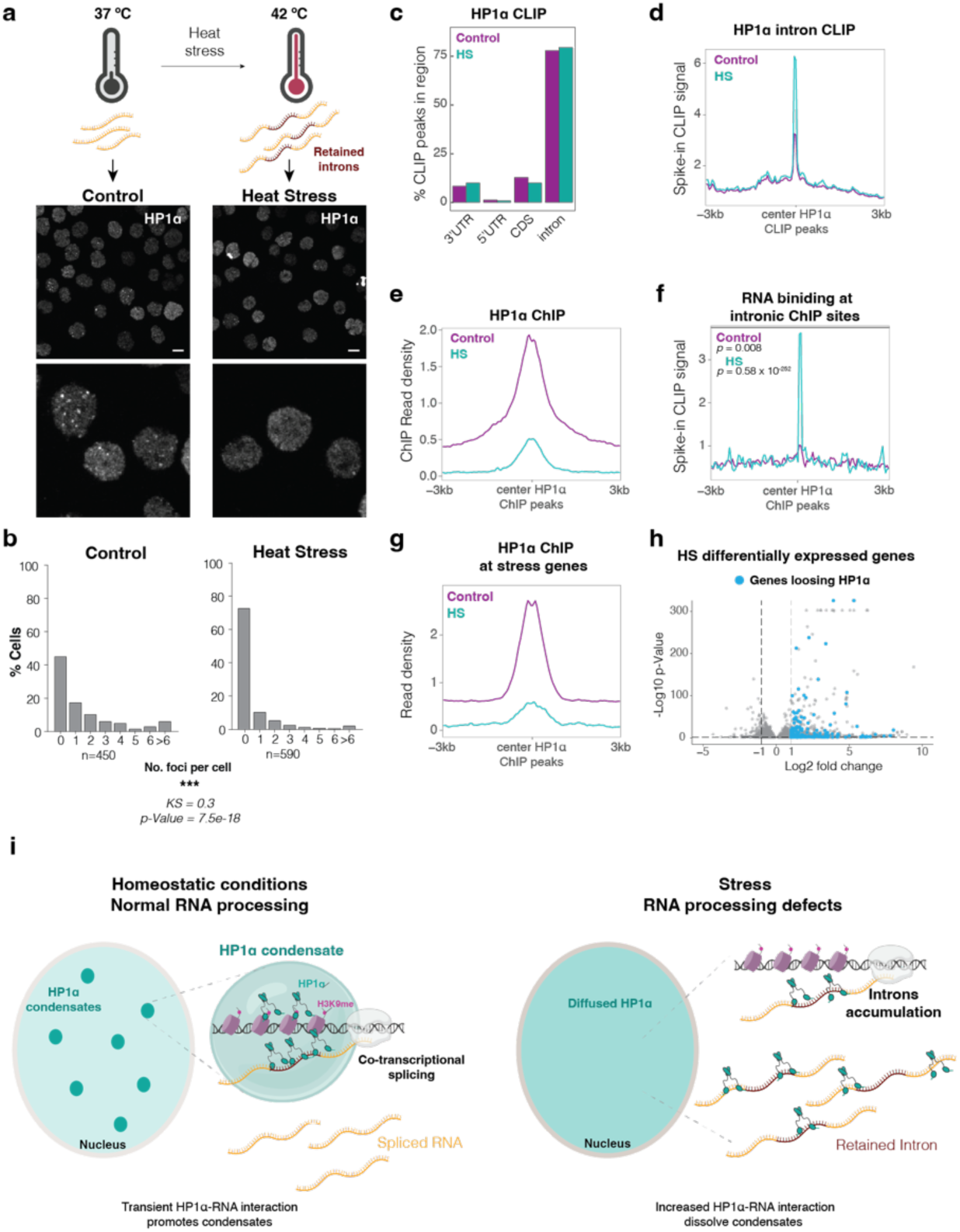
IR during stress regulates HP1α condensates. **a,** Top: schematic of the heat stress treatment. Bottom: maximum intensity projection confocal images of HP1α staining in K562 cells during stress. Magnified regions are shown below. Scale bar: 20 μm. **b,** Quantification of HP1α foci per cell. *n:* number of cells analyzed. KS statistic and *p*-value from the Kolmogorov-Smirnov test are shown. **c,** Relative distribution of HP1α CLIP-seq peaks across transcript features in control and HS cells. CDS: coding sequence. **d,** Metagene plots of HP1α spike-in normalized CLIP-seq signal in control and HS cells. **e,** HP1α ChIP-seq signal before and after heat stress across all HP1α genomic binding sites. **f,** Metagene plot showing HP1α–RNA binding enrichment at genomic intronic regions associated with HP1α. *p*-value from Pearson’s Chi-Square test of independence is indicated. **g**, HP1α ChIP-seq signal at stress-responsive genes in control and HS. **h,** Volcano plot of differentially expressed genes upon stress. Genes showing loss of HP1α ChIP signal are highlighted in blue. **i,** Model of the HP1α–intron regulatory loop. Left: under normal conditions, introns are co-transcriptionally spliced and HP1α–RNA interactions are restricted to chromatin, promoting HP1α–chromatin condensation. Right: upon stress or in disease, intron-containing transcripts accumulate in the nucleoplasm, leading to condensate dissolution and loss of silencing. Whether HP1α–RNA interactions act in cis or involve trans-acting RNAs remains to be determined.

Consistent with the localization KO mutants, genome-wide profiling of HP1α-RNA interactions confirms preferentially binding to introns (Fig. 5c). Spike-in normalized CLIP-seq reveals increased HP1α signal at intronic RNA sites following heat stress (Fig. 5d, Extended Data Fig. 5e), while ChIP-seq shows a global reduction in chromatin occupancy (Fig. 5e, Extended Data Fig. 5e), particularly at intronic regions (Extended Data Fig. 5f). Under stress, sharp ChIP peaks are replaced by a broader, lower-amplitude signal (Extended Data Fig. 5f), consistent with a transition from site-specific chromatin binding to a more diffuse binding. In addition, spike-in–normalized CLIP-seq reveals increased RNA association at HP1α ChIP sites where chromatin occupancy is reduced, consistent with a redistribution of HP1α from chromatin- to RNA-bound states (Fig. 5f, Extended Data Fig. 5e). Together, these data indicate that stress induces HP1α relocalization from chromatin to intron-containing transcripts in the nucleoplasm, resulting in condensate dissolution. This stress-induced changes recapitulate the phenotype of the HP1α localization mutants identified in our image-based CRISPR screen.

We then asked whether the loss of HP1α condensates functionally contributes to the stress response by relieving silencing and/or promoting RNA Polymerase II mobilization at stress-responsive genes. We first identified a subset of heat-responsive genes by RNA-seq (Extended Data Fig. 5g) and examined HP1α chromatin association at these loci. The observed loss of HP1α at stress-responsive genes (Fig.5g), including PKNOX1 and FRS2 (Extended Data Fig. 5e) suggests that HP1α relocalization may facilitate transcriptional changes in response to stress. Indeed, a substantial portion of stress genes (175/925), including heat shock proteins and chaperones, lose HP1α binding upon stress (Fig. 5h), underscoring the importance of HP1α in stress response. This functional role is further supported by the global differences in the transcriptional stress response observed in *HP1α* KO cells and in cells expressing the HP1α 6A mutant (Extended Data Fig. 5h), highlighting the contribution of HP1α and its Hinge domain in orchestrating gene expression changes during stress adaptation.

Overall, the loss of HP1α condensates under heat stress highlights the central role of HP1α-RNA interactions in maintaining heterochromatin structure and function. These findings, which are consistent with previous reports of heterochromatin loss under stress conditions^47^, provide a mechanistic basis for how IR contributes to stress adaptation, by altering HP1α localization and enabling the expression of protective genes.

Based on our results, we propose a model in which HP1α-intron interactions mediate a crosstalk between RNA processing and heterochromatin (Fig. 5i). In this model, H3K9me recruits HP1α to genomic regions, and RNA promotes condensate formation by lowering the critical concentration required to form condensates. Because pre-mRNAs are rapidly co-transcriptionally spliced, only a small fraction of RNA in proximity to chromatin contains introns^30–33^. As such, HP1α–intron interactions are transient, both spatially and temporally restricted at chromatin. During stress or pathological conditions in which intron-containing transcripts accumulate in the nucleoplasm, the concentration, composition, and subcellular localization of RNA are altered. This leads to increased HP1α–RNA interactions in the nucleoplasm, affecting chromatin interactions and resulting in the disassembly of HP1α condensates. We propose that RNA is not only a key structural component of heterochromatin but also a critical regulator that enables crosstalk between chromatin organization and RNA processing.

## Discussion

The canonical view of heterochromatin as static and transcriptionally silenced has been challenged^4,5,8,48^. The core heterochromatin protein HP1α is highly dynamic, showing rapid on-off rates within milliseconds, while forming spatially and functionally discrete condensates^4,5^. RNA is crucial component of heterochromatin that, when removed, alters heterochromatin architecture^3,10,11^.

To investigate the mechanisms underlying subcellular organization of heterochromatin compartments, we developed an image-based CRISPR screen for regulators of HP1α condensates. Our approach offers several advantages: (i) it uncovers novel regulatory pathways; (ii) it monitors endogenous HP1α protein, avoiding artifacts from over-expression and tagging^49^; (iii) it analyzes compartments in cells, preserving the cellular molecular composition; and (iv) it enables rapid imaging and scoring of condensates in millions of cells without the need for complex analysis pipelines. This high-throughput method can be applied to study other condensates and subcellular bodies, enabling to better understand the link between subcellular bodies structure and function.

We discovered that unspliced intronic RNA transcribed in heterochromatin regions regulates HP1α condensates by interacting co-transcriptionally with HP1α. These findings indicates that RNA and RNA processing can modulate HP1α localization, leading to several conceptual advances and a revised model of heterochromatin, as discussed below.

It is well established that RNA is a central component of nuclear condensates, such as nuclear speckles, Cajal bodies, and the nucleolus^50,51^. It serves as a scaffold for condensate assembly, and changes in RNA composition or localization regulate condensate formation and function^50^. Recent findings have highlighted RNA’s role in regulating transcriptional condensates through interactions with transcription factors^52,53^. Our findings extend these observations to heterochromatin, a nuclear compartment where the role of RNA may seem less intuitive. We propose that RNA acts as a key regulatory molecule, modulating interactions between chromatin and chromatin-associated factors in both active and repressed nuclear compartments, thereby providing a mechanistic model for the counterintuitive presence of RNA in heterochromatin. Furthermore, we propose that RNA processing enzymes regulate chromatin organization by fine-tuning RNA composition, localization, and concentration. The crosstalk we uncover between chromatin and RNA processing provides a molecular basis for the observed link between splicing and chromatin regulation^54,55^ and introduces a new perspective on genome regulation— positioning RNA processing as a surveillance system that not only shape the proteome but also governs the composition and localization of RNA across nuclear compartments.

HP1α localizes to interspersed heterochromatin within gene-rich regions—often associated with transposable elements— as well as to large heterochromatin domains at pericentromeres, telomers, and subtelomeres^56^. While RNA derived from pericentromeric repeats and transposons has been implicated in heterochromatin regulation^57^, our study focuses on non-repetitive regions, owing to the challenges associated with analyzing repeat-rich sequences. This leaves open important questions about the contribution of repetitive RNAs to HP1α function. Notably, introns are enriched in repetitive elements such as SINEs and LINEs and share sequence and structural features with repeat-derived RNAs^58,59^. Moreover, activation of both transposons and satellite repeats has been reported in response to cellular stress^60–63^. To explore this possibility, we analyzed our RNA-seq and CLIP-seq datasets using Classification of Ambivalent Sequences using *K*-mers (CASK)^64^. We observed that a subset of reads mapped to repetitive elements, including satellite repeats and various transposons (Extended Data Fig.5i, j), suggesting that repeat-derived RNAs may regulate HP1α through mechanisms akin to intronic RNAs, consistent with recent findings in the context of mouse major satellite repeats^65^.

Defects in RNA-mediated condensates contribute to pathogenesis of cancer, neurodegeneration, and infections^66^. The upregulation of stress protective genes with the loss of HP1α foci underscores the functional implication of HP1α organization in condensates, raising the possibility that changes in heterochromatin might represent a general mechanism underlying the pathogenesis and functional consequences of IR and splicing defects in diseases^67^. Supporting this hypothesis, loss of heterochromatin and HP1α foci has been observed in the cortices of mice affected by ALS, a neurodegenerative disease linked to IR^68^. This suggests that the heterochromatin-RNA processing axis may be a general mechanism in biology with broad implications in the pathobiology of processed linked to IR.

Lastly, contrary to traditional views, our work implies that mammalian heterochromatin can harbor actively transcribed genes that are marked by H3K9me3 and bound by HP1α, and that RNA contributes to the functional organization of heterochromatin in mammals. Our data, along with recent studies reporting transcription within inactive B compartments^40^, support a model in which heterochromatin is a dynamic compartment that can accommodate transcription and even rely on RNA for its structural or regulatory functions. In this model, HP1α is not a strict repressor, but rather a versatile effector whose function is shaped by RNA interactions, chromatin context, and cellular state.

## Acknowledgments

We thank A. Gitler, G. Narlikar, J. Gross, K. Daniels, J. Sage, N. Altemose and members of the Sanulli laboratory for helpful discussions and suggestions; N. Neff and the rest of the Stanford Chan Zuckerberg Biohub Center team for ongoing support. We thank James Medwid and Ashfeen Nawar for histone and hHP1α protein purifications. Mass Spectrometry was provided by the Mass Spectrometry Resource at UCSF (A.L. Burlingame, Director) supported by the Dr. Miriam and Sheldon G. Adelson Medical Research Foundation (AMRF). This work was supported by funds from the Chan Zuckerberg Biohub (to S.S.), the Emerson Collective Stanford Cancer Institute-Goldman Sachs Foundation Cancer Research Fund (to S.S), the Searle award (to S.S), and NIH DP2GM149752 (to S.S.). S.S. is a Chan Zuckerberg Biohub Investigator and holds a career award from the Kinship Foundation.

## Contributions

SS conceived and supervised the project. SS and RV performed the CRISPR screen. MMW designed and performed the majority of experiments including generating cell lines, imaging, RNA-seq, ChIP seq, bioinformatic analyses, cell culture assays, analyses of experimental data. SZ performed CLIP-seq, RNA fractionation, and long-read sequencing. CC analyzed AS, CLIP-seq data, and CASK. AW contributed to Punctatools data analysis. SS and MP performed *in vitro* phase assays and AR collected confocal images of multi-phases condensates. AA helped with cell cultures and confocal imaging, and performed ICE. CCC sorted cells for the screen. JT, DY, and KS generated the library of sgRNA. GLL performed PlaB treatments and IP. MJC performed MS. SS wrote the bulk of the manuscript with input from the authors. All authors discussed the results and commented on the manuscript.

## Inclusion & ethics statement

All collaborators of this study have fulfilled the criteria for authorship required by Nature Portfolio journals have been included as authors, as their participation was essential for the design and implementation of the study. Roles and responsibilities were agreed among collaborators ahead of the research. This work includes findings that are locally relevant, which have been determined in collaboration with local partners. This research was not severely restricted or prohibited in the setting of the researchers, and does not result in stigmatization, incrimination, discrimination or personal risk to participants. Local and regional research relevant to our study was taken into account in citations.

## Materials & Correspondence

Correspondence and requests for materials should be addressed to SS.

